# Human *CCL3L1* copy number variation, gene expression, and the role of the CCL3L1-CCR5 axis in lung function

**DOI:** 10.1101/249508

**Authors:** Adeolu B Adewoye, Nick Shrine, Linda Odenthal-Hesse, Samantha Welsh, Anders Malarstig, Scott Jelinsky, Iain Kilty, Martin D Tobin, Edward J Hollox, Louise V Wain

## Abstract

The CCL3L1-CCR5 signaling axis is important in a number of inflammatory responses, including macrophage function, and T-cell-dependent immune responses. Small molecule CCR5 antagonists exist, including the approved antiretroviral drug maraviroc, and therapeutic monoclonal antibodies are in development. Repositioning of drugs and targets into new disease areas can accelerate the availability of new therapies and substantially reduce costs. As it has been shown that drug targets with genetic evidence supporting their involvement in the disease are more likely to be successful in clinical development, using genetic association studies to identify new target repurposing opportunities could be fruitful. Here we investigate the potential of perturbation of the CCL3L1-CCR5 axis as treatment for respiratory disease. Europeans typically carry between 0 and 5 copies of *CCL3L1* and this multi-allelic variation is not detected by widely used genome-wide single nucleotide polymorphism studies. We directly measured the complex structural variation of *CCL3L1* using the Paralogue Ratio Test (PRT) and imputed (with validation) CCR5del32 genotypes in 5,000 individuals from UK Biobank, selected from the extremes of the lung function distribution, and analysed DNA and RNAseq data for *CCL3L1* from the 1000 Genomes Project. We confirmed the gene dosage effect of *CCL3L1* copy number on *CCL3L1* mRNA expression levels. We found no evidence for association of *CCL3L1* copy number or CCR5del32 genotype with lung function suggesting that repositioning CCR5 antagonists is unlikely to be successful for the treatment of airflow obstruction.

## Introduction

Genome-wide association studies have identified thousands of disease-gene associations leading to new disease insight and potential new approaches to treatment. It has been shown that drug targets supported by genetic studies have an increased chance of success in clinical development [1]. Even so, only a subset of candidate drugs will make it through to the clinic. Identifying opportunities for repositioning existing drugs and targets is therefore an appealing prospect and using genetic studies to define alternative indications for an already-approved drug is a promising approach.

The Mip1alpha (encoded by *CCL3* and *CCL3L1)-CCR5* signaling axis is important in a number of inflammatory responses, including macrophage function, and T-cell-dependent immune responses [2]. It is perturbed by CCR5 antagonists such as Pfizer’s maraviroc, the only CCR5 antagonist to be approved by the United States Food and Drug Administration [3, 4]. Identification of a genetic association of variants within the genes involved (*CCR5* and *CCL3/CCL3L1*) would strongly support the potential use of CCR5 antagonists in the treatment of respiratory conditions [5].

In mice, MIP1alpha is implicated in virus-mediated inflammation of the lung, pulmonary eosinophilia following paramyxovirus infection, clearance of pulmonary infections [6, 7], and in the response to respiratory syncytial virus infection [8-10]. In humans Mip1apha controls the recruitment of immune cells to inflammatory foci, and increased levels of Mip1alpha mRNA are found in bronchial epithelial cells of COPD patients [11], and increased protein levels in the sputum of COPD patients [12] where increased macrophage and neutrophil infiltration in the lung is a key pathology.

The *CCR5* gene in humans has a 32bp exonic deletion allele (rs333, CCR5d32) with a minor allele frequency of between 5-15% in Europeans [13]. This allele causes a translational frameshift and abrogates expression of the receptor at the cell surface, such that homozygotes for the deletion allele lack any functional CCR5 receptor [14, 15]. This variant has been strongly and repeatedly associated with resistance to HIV infection and slower HIV progression, as CCR5 is a common coreceptor for HIV entry into T-lymphocytes [16]. The CCR5d32 allele has been suggested to confer a reduced risk of asthma in children in one study [17] although this has not been replicated [18, 19].

In humans, there are two isoforms of Mip1alpha, the LD78a isoform encoded by the CCL3 gene and the LD78b isoform encoded by the paralogous *CCL3L1* gene [20, 21]. The two isoforms differ by three amino acids, but only one of these small changes, a serine to proline change at position 2 of the mature protein, alters the affinity to the cell surface receptor CCR5, with the beta isoform (*CCL3L1*) having approximately six-fold greater affinity [22] for CCR5 than the alpha isoform (*CCL3*).

The *CCL3L1* gene is part of a complex structurally variable region, although the *CCL3* gene is not. The *CCL3L1* gene and the neighboring *CCL4L1* gene are tandemly repeated with the total diploid copy number ranging from 0 copies to 6 copies in Europeans [23, 24]. Higher copy numbers are observed elsewhere, for example 10 in Tanzanians [25] and 14 in Ethiopians [26]. Previous studies have shown evidence of a gene dosage effect, with *CCL3L1* gene dose reflected in mRNA levels as well as in the ability to chemoattract monocytes [27, 28].

Measuring *CCL3L1* multiallelic copy number variation has been challenging [29]. Early studies used qPCR assays with a low signal:noise ratio [23, 30, 31], but assays based on the paralogue ratio test (PRT), allowed more accurate estimation of diploid copy number [24, 32]. Because of the challenges in measuring *CCL3L1* copy number in sufficiently large and well-powered sample sizes, the effect of structural variation of the genes encoding the Mip1alpha-CCR5 ligand-receptor pair has not been adequately explored.

In this study, we set out to confirm previous reports that *CCL3L1* copy number is associated with CCL3L1 gene expression, then measure *CCL3L1* copy number and *CCR5d32* genotype in 5000 individuals from UK Biobank, and finally test for association with lung function. Furthermore, we validated our copy number typing approach and observed copy number frequencies using publicly available sequence data from the 1000 Genomes Project. For *CCL3L1* copy number measurement in the 5000 individuals from UK Biobank, we used a triplex paralogue ratio test (PRT) which is considered to be the gold standard approach for measurement of this copy number variation [24, 29]. For genotyping of CCR5d32 in UK Biobank, we used a standard genotype imputation approach with additional PCR validation. We tested for association with extremes of Forced Expired Volume in 1 second (FEV1) as a binary trait. This study is the largest analysis of the effect of CCL3L1 copy number and CCR5d32 genotypes on lung function undertaken to date.

## Methods

### Sample selection

Individuals were selected from the UK BiLEVE [33, 34] subset of UK Biobank. In brief, 502,682 individuals were recruited to UK Biobank of whom 275,939 were of self-reported European-ancestry, and had two or more measures of Forced Expired Volume in 1s (FEV1) and Forced Vital Capacity (FVC) measures (Vitalograph Pneumotrac 6800, Buckingham, UK) passing ATS/ERS criteria [35]. Based on the highest available FEV1 measurement, 50,008 individuals with extreme low (n=10,002), near-average (n=10,000) and extreme high (n=5,002) % predicted FEV1 were selected from amongst never-smokers (total n=105,272) and heavy-smokers (mean 35 pack-years of smoking, total n=46,758), separately.. For this study, we selected 2500 age-matched European-ancestry heavy smokers from the extreme high and extreme low % predicted FEV1 subsets defined for the UK BiLEVE study (Figure 1, Table 1). DNA samples for these 5000 individuals were prepared by UK Biobank and provided back to the University of Leicester with new identification codes such that typing of *CCL3L1* copy number and *CCR5d32* was blinded to lung function status. Positive control samples for the copy number typing were from the Human Random Control panel from Public Health England.

**Figure 1.**
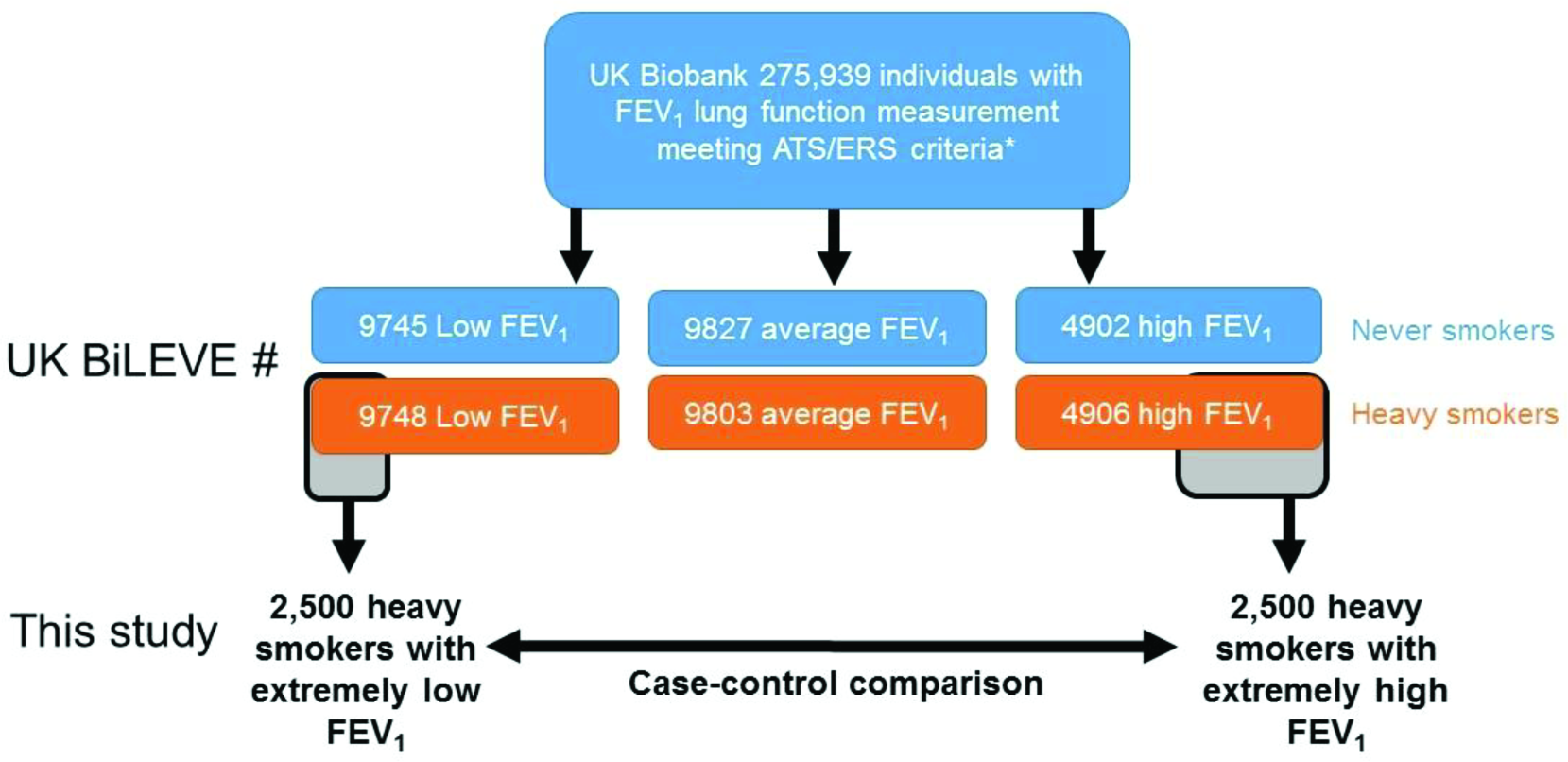
Study design. FEV_1_ is percent predicted FEV_1_. *Lung function measurement quality control defined previously [33] #Final numbers after quality control [33]

**Table 1.** Demographics of selected UK Biobank cohort.

|  | **Low FEV<sub>1</sub> (n=2500)** | **High FEV<sub>1</sub> (n=2500)** |
| --- | --- | --- |
| n (%) male | 1250 (50%) | 1250 (50%) |
| Age | 56.9 / 7.9<br>(40, 70) | 56.9 / 7.9<br>(40, 70) |
| Pack-years | 40.6 / 22.5<br>(10.8, 301.0) | 29.37 / 13.4<br>(10.5, 134.0) |
| Pack-years as a proportion of lifespan. | 0.96 / 0.47<br>(0.42, 7.00) | 0.70 / 0.29<br>(0.42, 3.03) |
| FEV<sub>1</sub> (litres) | 1.50 / 0.47<br>(0.36, 3.38) | 3.64 / 0.73<br>(2.02, 6.72) |
| Percent predicted FEV<sub>1</sub> | 51.4 / 11.0<br>(14.9, 74.5) | 123.3 / 8.2<br>(112.8, 205.7) |

Values are Mean / SD (range), unless stated.

### CCL3L1 copy number estimation in UK Biobank and 1000 Genomes Project samples using the paralogue ratio test (PRT)

*CCL3L1* copy number was determined using a triplex paralogue ratio test (PRT) assay as used previously [24, 26]. Briefly, PRT is a comparative PCR method that amplifies a test and reference locus using the same pair of primers, followed by capillary electrophoresis and quantification of the two products [32, 36]. The triplex assay produced three independent estimates of copy number per test, of which the average was taken as a representative copy number value. The three values were consistent in 95% of samples, however, for 5% of samples the value from the LTR61A PRT assay was significantly lower than the other two PRT values, and an average of the two consistent PRTs was taken in these 5% of samples. For each typing experiment, 4 positive controls of known copy number were also included, as previously [26, 37]. The copy number values clustered about integer copy numbers, and a Gaussian mixture model was fitted to allow assignment of individuals to an integer copy number call using CNVtools [38]. For the 5000 individuals from UK Biobank, 58 individuals were selected by UK Biobank investigators as blind spiked duplicates as part of the quality control check to ensure genotyping accuracy. Copy numbers from UK Biobank samples are available from UK Biobank at http://www.ukbiobank.ac.uk/data-showcase/.

### Gene Expression levels in 1000 genomes project lymphoblastoid cell lines

Matched RNAseq data that is publically available for the 1000 genomes samples were grouped based on *CCL3L1* copy number and analysed for their differential expression using Cufflinks v2.1.1 [39]. This allows measurement of the effect of genomic copy number of *CCL3L1* on gene expression levels. The analyses were all performed on ALICE High Performance Computing Facility at the University of Leicester. The RNAseq data were generated by (Lappalainen et al. 2013) and deposited in EBI ArrayExpress (accessions E-GEUV-1, E-GEUV-2, E-GEUV-3). Using Cufflinks, the fragments per kilobase of transcript per million fragments mapped (FPKM) values were estimated by applying a statistical model that normalises the mapped reads by length and their abundance. Briefly, the fragment reads are divided by transcript size and the total number of reads and then adjusted to 1 kb and 1 million reads.

### Genotyping of CCR5d32 polymorphism

Imputation to 1000 Genomes Project Phase 1+UK10K reference panel [40] and PCR were used to genotype the CCR5del32 polymorphism (rs333) in the 5000 UK Biobank individuals. Phasing and imputation were undertaken with SHAPEIT v2.r790 [41] and IMPUTE2 v2.3.1 [42]. For individuals with imputation posterior probability <0.95 (431 samples), and an additional 20 samples that were imputed as homozygous for the minor del32 allele, we validated the imputation results using direct PCR genotyping. Duplicates of a random selection of 28 of individuals were included as a quality control check for genotyping reproducibility (genotyping was also blinded to duplicate status). Genotypes from UK Biobank samples are available from UK Biobank at http://www.ukbiobank.ac.uk/data-showcase/.

### CCL3L1 copy number estimation from sequencing data for 1000 Genomes Project individuals

1000 genomes phase 3 whole genome aligned Bam files generated from Illumina platforms available from the European Bioinformatics Institute (ftp://ftp.1000genomes.ebi.ac.uk/vol1/ftp/data_collections/) were downloaded and the genomic region including *CCL3L1* (hg19:chr17:33670000-35670000) was analysed using CNVrd2 [43]. Using 500bp window sequence read depth, the sequence read depth was calculated across the region for all 2502 genomes from 26 populations, and standard deviation/quantile calculated for each window. The segmentation scores obtained from this analysis were clustered into different groups using a Gaussian mixture model. A prior information for all populations was estimated using the expectation maximisation (EM) algorithm on a population group with clear clusters of segmentation scores. The prior information (means, standard deviations and proportions of the mixture components) was fed into Bayesian model to infer *CCL3L1* integer copy number in all populations. Copy number estimates are available from dbVar (https://www.ncbi.nlm.nih.gov/dbvar) under study accession number nstd155.

### Association analysis

We tested for association of *CCL3L1* copy number and CCR5d32 genotype separately with lung function extremes (as a binary trait) using logistic regression with pack-years of smoking and the first ten principal components (obtained previously using full genome-wide SNP genotyping data to adjust for fine-scale population structure as covariates [33]. For CCR5d32, a genotypic genetic model was assumed for the primary analysis. We then fitted a full linear regression model that included CCR5d32 genotype (genotypic mode), *CCL3L1* copy number, pack years, 10 principal components and a term for the interaction of *CCR5d32* and *CCL3L1*.

## Results

Using CNVrd2, we typed *CCL3L1* copy number from whole genome sequence alignments for 2502 individuals from the 1000 Genomes project (Figure 2a). The data were grouped into large superpopulations, as defined by the 1000 Genomes Project [44], and our analysis confirmed previous observations that Europeans have the lowest *CCL3L1* diploid copy number, ranging between 0 and 5 with a mean copy number of 1.97, and sub-Saharan Africans have the highest diploid copy number, ranging between 1 and 9 with a mean of 4.19, which is more than twice as high as Europeans (Table 2)[24, 25].

**Figure 2.**
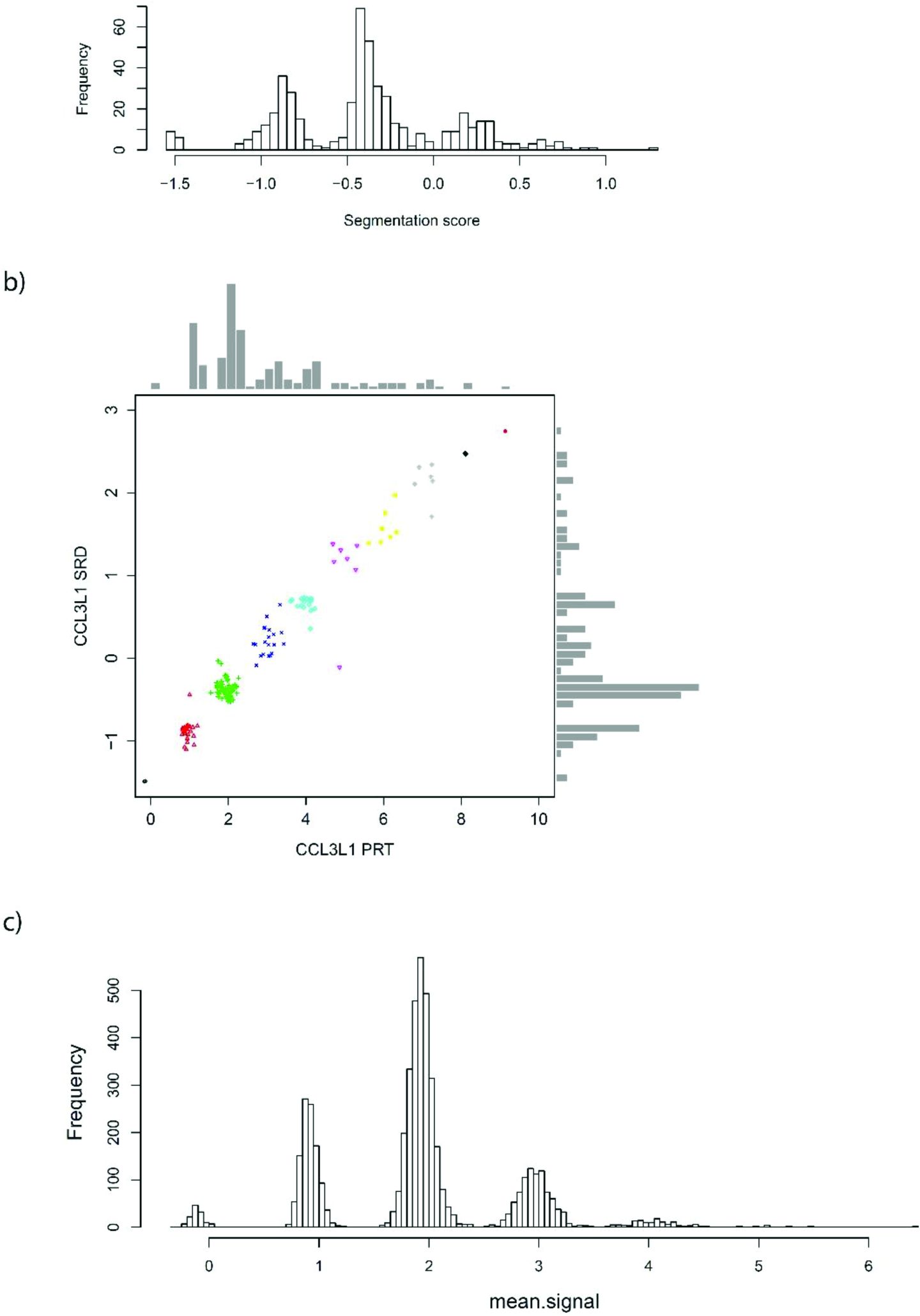
*CCL3L1* Copy number typing. a) Histogram of raw copy number estimates of 1000 Genomes Project samples from sequence read depth represented as segmentation scores on the x axis, generated by CNVrd2, with higher scores reflecting higher copy number. b) Validation of 144 1000 Genomes Project samples using PRT (x axis) against estimates made from sequence read depth. Colours/symbols in the scatterplot represent different integer copy numbers inferred from PRT clusters. c) Histogram of raw copy number estimates using PRT for the UK Biobank cohort.

**Table 2.** CCL3L1 copy number frequency distributions in 1000 Genomes data.

| superpopulation | n | average<br>copy<br>number | minimum<br>copy<br>number | maximum<br>copy<br>number |
| --- | --- | --- | --- | --- |
| **AFR (Sub-Saharan African)** | 661 | 4.19 | 1 | 9 |
| **AMR (Admixed American)** | 347 | 2.71 | 0 | 8 |
| **EAS (East Asian)** | 504 | 3.52 | 0 | 9 |
| **EUR (European)** | 501 | 1.97 | 0 | 5 |
| **SAS (South Asian)** | 489 | 2.39 | 0 | 7 |

For 144 individuals from the CEU (n=96) and YRI (n=48) populations of the 1000 Genomes project, we also determined *CCL3L1* copy number using the PRT approach (Figure 2b). There was strong concordance between results, with discrete clusters of raw data, representing individual integer copy numbers, formed, particularly at low copy number. For the range seen in Europeans (copy numbers 0 to 5), there are seven clear discrepancies which gives an joint error rate of 5%.

To confirm previous studies that reported an association between *CCL3L1* copy number and *CCL3L1* mRNA levels, we compared the 1000 Genomes Project *CCL3L1* copy numbers with transcript levels of *CCL3L1* and its non-copy number variable paralogue *CCL3*, as generated by RNAseq of the corresponding B-lymphoblastoid cell lines (Figures 3a, 3b). Comparison with transcript level estimates using RNAseq data showed a clear positive correlation between *CCL3L1* copy number and expression level (figure 3b, r^2^=0.25, p<2x10^-16^). We used the specific sequence changes between *CCL3L* and *CCL3* to distinguish transcripts from either gene, and confirmed this by showing that *CCL3* expression has no relationship with *CCL3L1* copy number (figure 3a, r^2^=0.006, p=0.087), as well as showing that individuals with zero copies of *CCL3L1* show no transcripts from *CCL3L1* (figure 3b).

**Figure 3.**
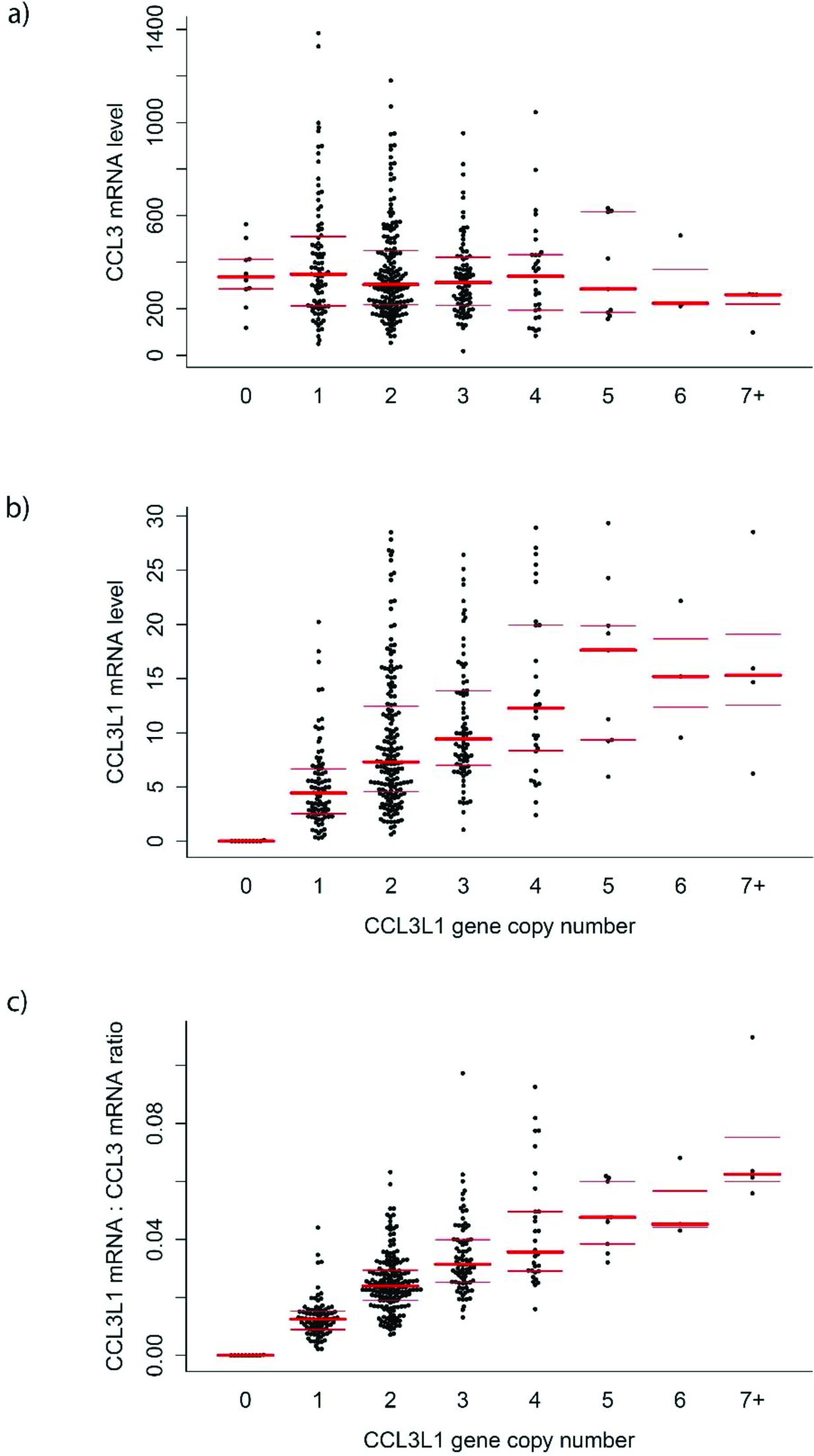
Copy number and expression level of *CCL3L1* and *CCL3* in lymphoblastoid cell lines. a) *CCL3* mRNA level (FPKM units) across different *CCL3L1* copy numbers. b) *CCL3L1* mRNA level (FPKM units) across different *CCL3L1* copy numbers. c) *CCL3L1.CCL3* mRNA ratio across different *CCL3L1* copy numbers. Individual data points are shown, with red bars indicating median and interquartile ranges.

We confirmed an increase of one to two orders of magnitude for *CCL3* transcript levels compared to *CCL3L1* transcript levels in B-lymphoblast cells. Following normalization of the *CCL3L1* expression levels to CCL3 expression levels, we show that *CCL3L1* transcript levels are closely correlated with gene copy number (Figure 3c, r^2^=0.5, p<2x10^-16^).

Having confirmed a relationship between gene copy number and transcript levels of *CCL3L1*, we investigated the relationships between CCL3L1 copy numbers, CCR5d32 genotype and lung function in individuals selected from the extremes of the lung function distribution in UK Biobank. We typed 5000 UK Biobank samples using PRT, with 19 failures. The results showed a clear mixture of Gaussian distributions centered on each integer copy number (Figure 2c). All 58 duplicates were consistently typed, resulting in an error rate between 0% and 4.7%. We observed clear distances between the clusters, further suggesting that the measurement error rate for this cohort is likely to be low.

We estimated *CCL3L1* integer copy numbers in all the samples using Gaussian mixture modelling (Table 3). The copy number range was consistent with previous observations in UK population [24], and with our estimation from the 1000 Genomes project samples. The two copy genotype was the most frequent with a frequency of 0.563. The *CCL3L1* zero copy null genotype is uncommon, with a frequency of 2.5% in the UK. 4993 of the 5000 UK Biobank samples were genotyped for CCR5d32 by imputation with the genotypes for 474 individuals validated using direct PCR analysis. There was no evidence that the genotype frequencies departed from Hardy-Weinberg equilibrium (chi-squared test, p=0.35) and the observed CCR5d32 deletion allele frequency was 0.11, consistent with previous estimates [13].

**Table 3.** *CCL3L1* copy number counts in UK Biobank data.

| CCL3L1 diploid copy number | Number of samples | Frequency |
| --- | --- | --- |
| 0 | 127 | 0.025 |
| 1 | 1046 | 0.210 |
| 2 | 2806 | 0.563 |
| 3 | 853 | 0.171 |
| 4 | 128 | 0.026 |
| 5 | 21 | 0.004 |
| Sum | 4981 | 0.999 |

A total of 4975 UK Biobank individuals had both *CCL3L1* copy number and CCR5d32 genotypes measured (2486 high and 2489 low FEV1, Table 4). There was no evidence of an association between *CCL3L1* copy number and CCR5d32 genotype (chi-squared test p=0.803).

**Table 4.** CCR5d32 genotype counts by CCL3L1 copy number in UK Biobank data.

| CCL3L1 copy number | CCR5d32 genotype |  |  |
| --- | --- | --- | --- |
|  | ref/ref | del32/ref | del32/del32 |
| 0 | 92 | 33 | 2 |
| 1 | 826 | 203 | 16 |
| 2 | 2197 | 574 | 31 |
| 3 | 662 | 181 | 9 |
| 4 | 99 | 28 | 1 |
| 5 | 15 | 6 | 0 |
| Sum | 3891 | 1025 | 59 |

We fitted a full model with both CCR5 genotypes (genotypic model) and *CCL3L1* copy number and an interaction term as described above. This was undertaken in order to identify whether particular combinations of *CCL3L1* copy number and CCR5d32 genotype were differentially associated with lung function. Pack years of smoking and 10 principal components were included as covariates. No associations were significant (Table 5).

**Table 5.**
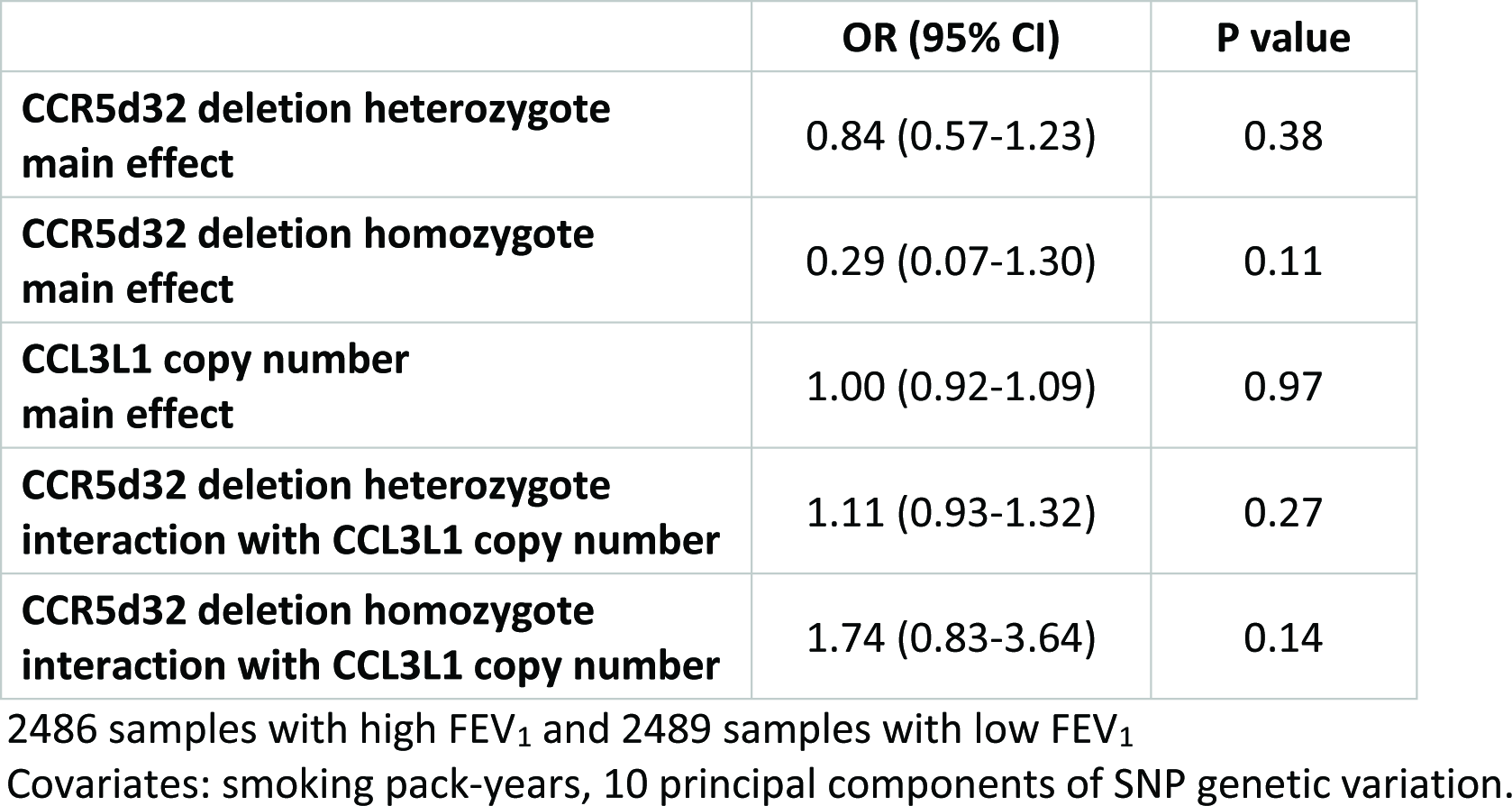
Association analysis of *CCR5* genotype and *CCL3L1* copy number with high vs low FEV_1_.

## Discussion

Our study provides robust large-scale confirmation of a gene dosage effect of CCL3L1 copy number on CCL3L1 mRNA levels, and also emphasises the strong dependence of *CCL3L1*:CCL3 mRNA ratio on copy number, with *CCL3L1* copy number accounting for 50% of total variation. Although it is clear that *CCL3L1* is expressed at much lower levels than *CCL3*, the MIP1alpha isoform encoded by *CCL3L1* (LD78beta) has a much stronger affinity to the CCR5 receptor than MIP1alpha isoform *CCL3* (LD78alpha). It therefore seems likely that the *CCL3L1* copy number variation mediates a biological effect *in vivo*. It should be noted that the expression data are from transformed B-lymphoblastoid cell lines, but a gene dosage effect is consistent with a study using fresh monocytes from 55 different individuals stimulated with bacterial lipopolysccharide [28].

Our analysis provides evidence that there is no effect of either *CCL3L1* copy number or CCR5d32 genotype, or any combinations of genotypes at the two loci, on lung function. This suggests that, although the Mip1alpha-CCR5 signaling axis can be disrupted by artificial CCR5 antagonists, there is no evidence that this axis has a functional effect on lung function and that development of new drugs to target this axis, or repurposing of existing drugs, might be of little or no therapeutic benefit in treating COPD.

We analysed approximately 5000 individuals. Whilst this represents a large sample size for labour-intensive PRT assays, it is a modest sample size in comparison with those employed in GWAS. That said, power was boosted by selecting from the extremes of the lung function distribution in the very large (n~500K) UK Biobank.

We reported PRT error rates of 2.5% for the 144 1000 Genomes Project samples and between 0% and 4.75% for the 4981 UK Biobank participants. A previous study using this PRT approach estimated an error rate of less than 0.1% [24], which suggests that much of the joint error rate for the PRT and sequence read depth could be due to errors in the sequence read depth approach.

The exact boundaries of the *CCL3L1* CNV have yet to be determined with precision but it is known to include the *CCL4L1* gene which encodes MIP1β [24]. The human genome assembly GRCh38 shows a single copy *CCL3L1/CCL4L1* repeat unit, and also includes the *TBC1D3* gene, encoding TBC1 Domain Family Member 3 [45-47]. The GRCh38 alternative assembly chr17_KI270909v1_alt shows two repeat units, both including *TBC1D3*. However an earlier assembly shows a complete contig with two repeat units carrying *CCL3L1/CCL4L1*, only one of which carries *TBC1D3*. ArrayCGH and fiber-FISH both confirm this is real heterogeneity by showing that the *TBC1D3* gene is included in some, but not all, tandemly repeated units in some individuals, together with *CCL3L1* and *CCL4L1* [26, 48]. Throughout this paper, and in most of the literature, *CCL3L1* CNV is used as a shorthand to describe the CNV of this complex repeat unit.

Given the gene content of this repeat unit, we would expect a gene dosage effect for *CCL4L1* and *TBC1D3*, in addition to *CCL3L1*, but this has not yet been confirmed. Our data do, however, show no effect of *CCL3L1* copy number on expression levels of its close paralogue, *CCL3*, which is immediately proximal to the CNV. This difference shows that the considerable variation in genome structure distal to the *CCL3* gene does not affect overall levels of *CCL3* expression.

In summary, we selected individuals from the extremes of the lung function distribution of a very large general population cohort. We found no association of *CCL3L1* copy number, nor of the CCR5d32 variant with lung function, as defined by FEV_1_.

## Acknowledgements

This research has been conducted using the UK Biobank Resource under Application Number 7140. This work was supported by funding from Pfizer, Inc to MDT, LVW and EJH. This article presents independent research funded partially by the National Institute for Health Research (NIHR). The views expressed are those of the author(s) and not necessarily those of the NHS, the NIHR or the Department of Health. Single nucleotide polymorphism genotyping of the UK BiLEVE subset of UK Biobank was funded by a Medical Research Council (MRC) strategic award to MDT, IPH, DPS and LVW (MC_PC_12010) (UK Biobank Application Number 648). LVW holds a GSK / British Lung Foundation Chair in Respiratory Research. MDT holds a Wellcome Trust Investigator Award. This research used the ALICE and SPECTRE High Performance Computing Facilities at the University of Leicester. We would like to thank Gurdeep Lall and Amelia Veselis for technical support.

